# Detecting Genes Associated with Pathogenicity and Antimicrobial Resistance in Three New Zealand Waterways

**DOI:** 10.1101/2020.04.16.045633

**Authors:** Meredith T. Davis, Anne C. Midwinter, Russell G. Death, Richard Cosgrove, Richard C. Winkworth

**Author notes:** Corresponding Author: Meredith T. Davis, School of Agriculture and the Environment, Massey University, Private Bag 11 222, Palmerston North, 4442, New Zealand.

## Abstract

**Background:** More than 100 zoonoses may be transmitted via water, among them enteric diseases are leading causes of human mortality. Traditional monitoring for zoonoses relies on culturing of indicator species, but environmental DNA (eDNA) provides an alternative, allowing direct testing for genetic loci associated with pathogenicity and/or antimicrobial resistance in zoonotic bacteria.

**Objectives:** To evaluate whether genes associated with Shiga toxin producing *Escherichia coli* (STEC) and antimicrobial resistance can be monitored in waterways using culture-free sampling of eDNA combined with PCR-based testing.

**Methods:** Water and sediment samples were collected from two sites on each of three rivers in Canterbury, New Zealand; sample sites were situated above and below reaches bordered by intensive dairy farming. Samples from each site were tested for genes typically associated with *Escherichia coli*, STEC serogroups O26 and O157, human pathogenic strains of STEC, and resistance to a broad range of antibiotics.

**Results:** Both culturing and genetic testing confirmed the presence of *E. coli* in all samples. In contrast, presence of genes associated with STEC and antibiotic resistance varied by season and substrate. The O157 serogroup was identified at more than twice as many sites as O26, with the latter more common in autumn samples. In autumn, genes associated with pathogenic STEC were detected in one Ashley and both Rangitata River samples but were present in all spring samples, except one Ashley and one Selwyn River collection. The antibiotic resistance gene was only identified in spring, predominantly at sites downstream of intensive dairying.

**Discussion:** While our sample is small this study indicates that genetic testing of eDNA can be a useful tool for monitoring the presence and persistence of zoonoses in waterways. How the presence of these genetic elements is related to that of pathogenic STEC and incidence of disease in humans now needs to be examined.

## Introduction

Globally, the degradation of freshwater ecosystems due to anthropogenic land use changes has been linked to outbreaks of waterborne disease in humans (OECD 2017). Industrialisation (Closs et al. 2016; Ratha 2019), urbanisation (Merritt et al. 2010; Power et al. 2018), and intensive agriculture (Clapcott et al. 2012; Gluckman 2017; Joy 2015) all have the potential to negatively affect freshwater ecosystems through removal of riparian vegetation (Hickey and Doran 2004; Sweeney and Newbold 2014), heavy metal accumulation (Hickey and Clements 1998), increased sediment and nutrient inputs (Waters 1995; Yan et al. 2016), and contamination with faecal effluent (Dangendorf 2004). In New Zealand rapid changes in land use (Julian et al. 2017), the degradation of freshwater ecosystems (Clapcott et al. 2012), and steady increases in outbreaks of notifiable bacterial zoonoses are all well documented (Ministry for the Environment and Statistics New Zealand 2017).

Enteric diseases, which may be waterborne, are a leading cause of human deaths worldwide, especially in children under five years of age in lower income countries (World Health Organization 2019). Currently more than 100 zoonoses are recognised as being transmitted in aquatic ecosystems. These include members of *Leptospira, Campylobacter, Escherichia*, and *Salmonella* (Fang 2014; Gluckman 2017; Shaw et al. 2016). Human and animal faecal effluent are primary sources of the microorganisms responsible for waterborne zoonoses (Reddy et al. 1981). In broad terms, the risk of contracting bacterial zoonoses from effluent contaminated waterways increases for those swimming in (McBride et al. 2002), gathering food from (Perkins et al. 2016; Rose et al. 2001) or eating undercooked produce irrigated with (Adator et al. 2018; King et al. 2012; Solomon et al. 2002) effluent contaminated water. However, in specific terms, the risks associated with using effluent contaminated recreational waterways remain poorly understood because they are complex, dynamic and multi-factorial (Colford Jr et al. 2007; Prieto et al. 2001).

Shiga toxin producing *Escherichia coli* (STEC) is an emerging group of zoonoses linked to ruminants (Colford Jr et al. 2007; Oporto et al. 2019) that may occur in in recreational waters and to be transmitted via the oral-faecal route (Swaggerty et al. 2018), STEC is the fourth most commonly reported zoonosis in the EU (Severi et al. 2016) and USA (European Center for Disease Prevention and Control 2018) and an emerging issue in New Zealand (ESR 2019). In humans STEC infections typically present as enteric disease that in severe cases may progress to haemolytic uraemic syndrome (HUS) or kidney failure (Karmali 2018). Strains of *E. coli* recognised as belonging to the STEC group are generally characterised by the presence of one or a pair of Shiga toxin producing genes (*stx1* and *stx2*) (Donnenberg et al. 1997; Perna et al. 1998). However, these strains are otherwise diverse; belonging to 129 serogroups and with more than 260 antigen combinations (Valilis et al. 2018). Typically, human pathogenic STEC strains contain the locus of enterocyte and effacement (LEE) island (Paton and Paton 1998) and, until recently, were most likely to be identified as belonging to the O157 serogroup (Centers for Disease Control and Prevention 2018; European Center for Disease Prevention and Control 2018). In New Zealand human cases of STEC disease tripled between 2014 and 2017 (Health and Environment Group ESR 2019). However, over this same period the proportion of cases linked to the O157 serogroup fell from 88.1% to 37.1% (Health and Environment Group ESR 2019). Non-O157 STECs (e.g., O26, O45, O111) are increasingly being linked to disease in humans (Germinario et al. 2016; Gill et al. 2019; Luna-Gierke et al. 2014; Severi et al. 2016). This observation may simply reflect improved screening (Health and Environment Group ESR 2019) or may be due to changes in serotype prevalence.

Culture for human pathogenic STEC involves the isolation of candidate organisms based on their ability to metabolise different carbohydrates (Amézquita-López et al. 2018) which is then followed by molecular characterisation. Molecular characterisation of STEC typically focuses on genes associated with virulence (i.e., *stx1* and *stx2, eae* located on the LEE island) and the serotypes most commonly found in human cases (e.g., *rfbE* for O157, *wzy* for O26) (Anklam et al. 2012; Franz et al. 2007). However, in most studies of environmental samples, the microbial communities are enriched, using growth media, for 18-24 hours prior to plating and identification (De Boer and Heuvelink 2000; Irshad et al. 2016). Enriching samples in this way is not ideal as it alters the composition of the original microbial community; effects may be due to contrasting media preferences (Amagliani et al. 2018) or interactions between community members (Chekabab et al. 2013; Mauro et al. 2013). At worst such changes may result in failure to detect potentially pathogenic community members.

Use of traditional culturing is further complicated in the case of STEC as strains may lose genes associated with virulence during incubation for enrichment or culturing (Senthakumaran et al. 2018; Tarr et al. 2019). An alternative to the traditional approach is to test DNA extracted directly from environmental samples for the presence of genes associated with virulence or antibiotic resistance (Bélanger et al. 2002; Werber et al. 2002). Testing of environmental DNA (eDNA) can provide a rapid and inexpensive survey of human pathogens at a site. This approach could be used to examine whether genes typically associated with pathogenic STEC (e.g., *stx1, stx2* and *eae*) are present at a given site. Although this approach does not identify pathogenic isolates, the presence of virulence genes in a given sample should raise concerns about the potential level of faecal contamination at that site. This approach allows us to identify sites where further investigation, including the isolation and testing of individual microbes, is warranted.

In this study we use a culture free, eDNA method to test for genes associated with STEC from sites on three Canterbury, New Zealand rivers. Samples were collected to coincide with seasonal peaks in human STEC cases during autumn and spring (Health and Environment Group ESR 2019). We tested both benthic sediments and water column samples for five genes frequently associated with human pathogenic STEC and the recurrently co-morbid, group 1 CTX-M β-lactamases which confer resistance to multiple antibiotics (Ishii et al. 2005; Valat et al. 2012). For each sample we also performed traditional *E. coli* counts to determine whether there was any relationship between colony numbers and the prevalence of these genes.

## Materials and Methods

### Sample Collection

Water and sediment samples were collected from the Ashley, Rangitata, and Selwyn rivers once in austral autumn and once in the spring of 2018. Collections were made at two sites along each river; these sites were 10-15 km apart with one above and the other below reaches bordered by high densities of intensive dairying (Fig.1). At each site, water and benthic sediment samples were collected into separate sterile containers. Three 1 L water and three 25 g sediment samples were collected at each site. Samples were packed on ice, transported to the laboratory, and processed within 24 hours of collection.

**Figure 1.**
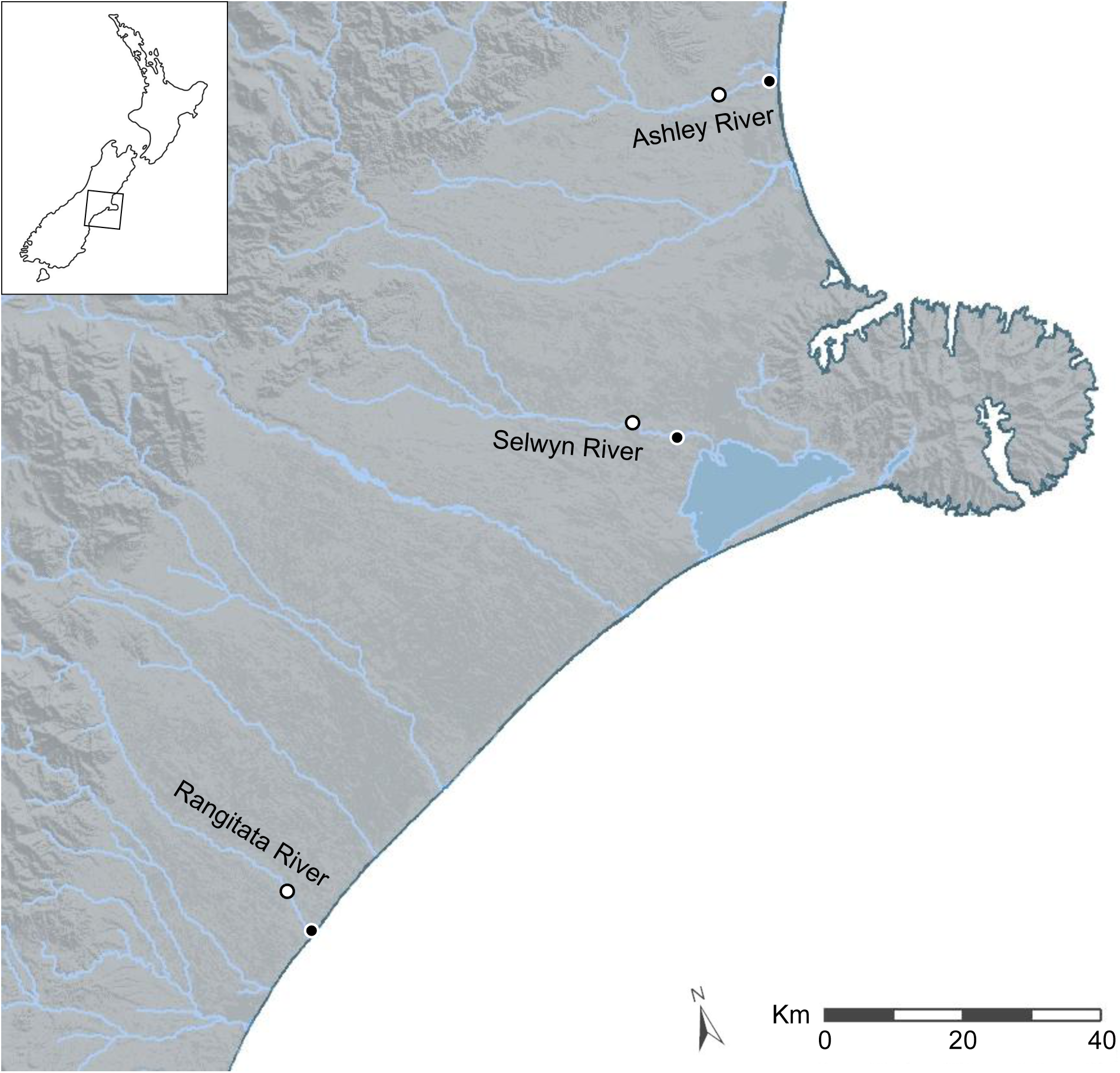
Map of central and southern Canterbury, New Zealand with the Ashley, Rangitata, and Selwyn rivers labeled. Locations at which water and sediment were collected during May and September 2018 are marked with circles; for each river the white centered dot indicates the above intensive dairy site and the black centered dot the below intensive dairy site.

### Sample Processing for Bacterial Culturing

Water column sample aliquots were diluted 1:10, 1:5, and 1:2.5 with sterile MilliQ H_2_O to a final volume of 50 ml. At sites where the level of suspended sediments was high, an additional 1:50 dilution was also prepared to ensure ease of enumeration. For each sample we prepared three technical replicates at each dilution (i.e., 9-12 dilutions per sample).

For sediment samples, 2 g of wet sediment was first transferred to a 5 ml microtube, 3 ml MilliQ H_2_O added, and the mixture then agitated vigorously for 30 seconds. Aliquots of the resulting supernatant were diluted 1:500, 1:200, 1:100, and 1:50 with MilliQ H_2_O to a final volume of 50 ml. For each sample we prepared three technical replicates at each dilution (i.e., 12 dilutions per sample).

For each dilution the total 50 ml volume was vacuum filtered through a single sterile 0.45 µm cellulose ester membrane filter (Merck KGaA, Darmstadt, Germany).

### Bacterial Culturing

Bacterial culturing followed United States Environmental Protection Agency method 1603 (EPA 2002). Each filter was placed onto a Difco Modified mTEC Agar (VWR, Radnor, PA) plate, incubated at 37.5 °C for two hours, and then incubated at 45 °C for 18-20 hours. Following incubation colonies resembling *E. coli* (red/magenta colonies) were counted.

### Sample Processing for Molecular Testing

Three 500 ml aliquots of water from each site were vacuum filtered through separate 0.45 µm cellulose ester membrane filters. Environmental DNA was extracted from half of each filter using the NucleoSpin® soil kit (Machery-Nagel GmbH and Co. KG, Düren, Germany) following the manufacturer’s instructions. For each sediment sample eDNA was extracted from three 0.5 g aliquots of wet sediment, using the NucleoSpin® soil kit.

### Molecular Testing for Specific Genes

The presence of five genes most commonly associated with human pathogenic STEC was evaluated using a polymerase chain reaction (PCR). Specifically, we targeted genes associated with serogroup specific antigen biosynthesis, *rfbE* for O157 and *wzy* for O26, the *stx*_1_ and *stx*_2_ toxin genes, and the intimate attachment and effacing gene, *eae*, using the primers reported by Anklam et al. (2012). We also investigated the presence of antimicrobial resistance by targeting the *bla* gene associated with group 1 CTX-M β-lactamases using the primers reported by Lalzampuia et al. (2013). Finally, as an amplification control we targeted the beta-glucuronidase gene, *uidA*, which is present in most *E. coli* (Anklam et al. 2012).

Amplification reactions were performed in 20 μl reaction volumes containing 0.5 × iQ PerfeCTa® qPCR ToughMix™, ROX™ (QIAGEN, Düsseldorf, Germany), 1 pM of each primer, and 2.5 μl of DNA template. Thermocycling was performed in a T1 thermocycler (Biometra GmbH, Göttingen, Germany) using standard cycling conditions including an initial denaturation at 94°C for 3 mins, followed by 35 cycles of 94°C for 30 s, 60°C for 30 s and 72°C for 1 min, with a final extension at 72°C for 5 mins. Amplification products were visualised using SYBR Safe (ThermoFisher Scientific, Waltham, MA) following electrophoresis on 2% Tris-acetate-ethylenediamine tetraacetic acid agarose gels.

## Results

### Bacterial Culturing

With just one exception, the spring sampling at the above intensive dairy site on the Selwyn River, *E. coli* counts were consistently higher from sediment samples than water column samples (Table 1). Moreover, counts were also higher for all but one sample taken at below intensive dairy sites. Only the spring sediment sample from the above intensive dairy site on the Rangitata River had a higher *E. coli* count than the corresponding sample from the below intensive dairy site.

**Table 1.**
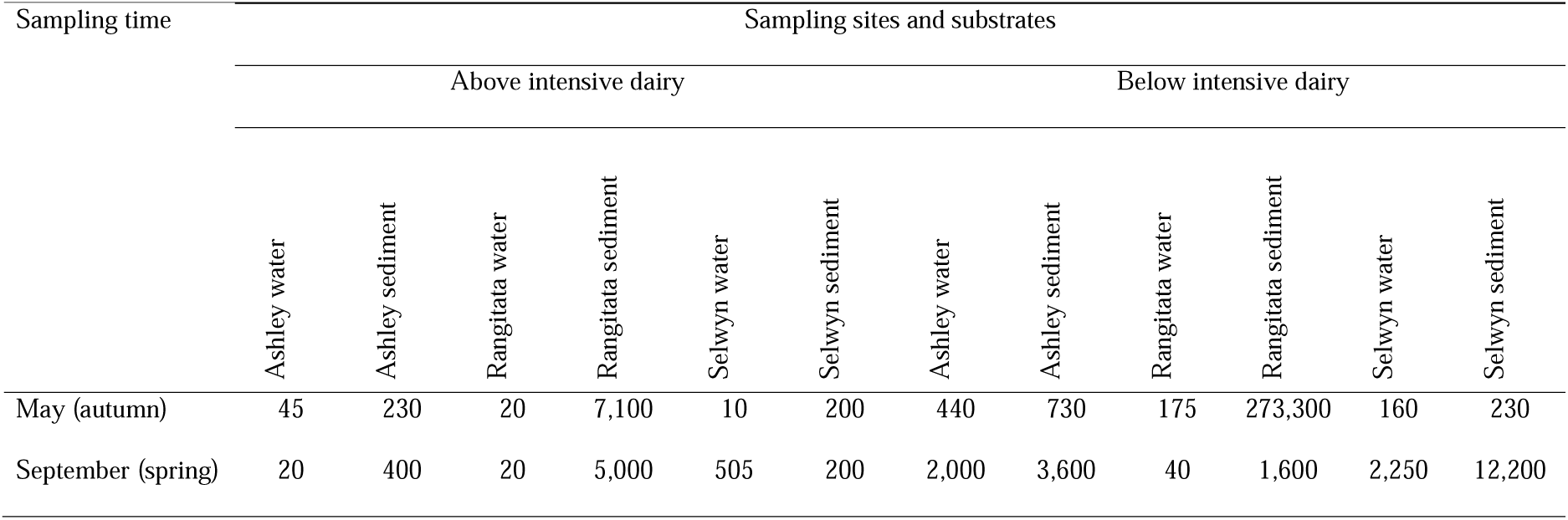
*Escherichia coli* colony counts, in CFU/100 ml, for water and sediment samples collected from the Selwyn, Rangitata, and Ashley rivers during May and September 2018.

| Sampling time | Sampling sites and substrates |  |  |  |  |  |  |  |  |  |  |  |
| --- | --- | --- | --- | --- | --- | --- | --- | --- | --- | --- | --- | --- |
|  | Above intensive dairy |  |  |  |  |  | Below intensive dairy |  |  |  |  |  |
|  | Ashley water | Ashley sediment | Rangitata water | Rangitata sediment | Selwyn water | Selwyn sediment | Ashley water | Ashley sediment | Rangitata water | Rangitata sediment | Selwyn water | Selwyn sediment |
| May (autumn) | 45 | 230 | 20 | 7,100 | 10 | 200 | 440 | 730 | 175 | 273,300 | 160 | 230 |
| September (spring) | 20 | 400 | 20 | 5,000 | 505 | 200 | 2,000 | 3,600 | 40 | 1,600 | 2,250 | 12,200 |

### Molecular Testing for Specific Genes

The *uidA* locus was successfully amplified from every sample in this study (Table 2). In contrast, detection of the three gene loci associated with pathogenic STEC (*stx1, stx2* and *eae*) and the gene associated with antibiotic resistance (*bla* CTX-M) varied with location, time, and substrate. The *stx1, stx2* and *eae* genes were more frequently detected in water samples in the autumn but in sediment samples in the spring. At both sampling times these three virulence genes were found together with the O26 (*wzy*) and O157 (*rfbE*) genes. The O26 marker was more frequently recovered from water, while that for O157 was evenly distributed between the substrates. The antibiotic resistance gene was detected in both substrates in samples from the Selwyn and Rangitata rivers.

**Table 2.** Presence of *Escherichia coli* control (*uidA*), STEC virulence (*stx1, stx2, eae*), serogroup (O26 *wzy* and O157 *rfbE*), and antibiotic resistance (*bla* CTX-M) genes in water and sediment samples collected from the Selwyn, Rangitata, and Ashley rivers during May and September 2018.

| Gene locus | Sampling sites and substrates |  |  |  |  |  |  |  |  |  |  |  |
| --- | --- | --- | --- | --- | --- | --- | --- | --- | --- | --- | --- | --- |
|  | Above intensive dairy |  |  |  |  |  | Below intensive dairy |  |  |  |  |  |
|  | Ashley water | Ashley sediment | Rangitata water | Rangitata sediment | Selwyn water | Selwyn sediment | Ashley water | Ashley sediment | Rangitata water | Rangitata sediment | Selwyn water | Selwyn sediment |
| May (autumn) |  |  |  |  |  |  |  |  |  |  |  |  |
| *uidA* | + | + | + | + | + | + | + | + | + | + | + | + |
| *stx1* | – | – | – | – | – | – | + | + | + | – | – | – |
| *stx2* | – | – | – | – | – | – | – | + | + | – | – | – |
| *eae* | – | – | – | – | – | – | + | + | + | – | – | – |
| *wzy* O26 | – | – | – | – | – | – | + | + | + | – | – | – |
| *rfbE* O157 | – | – | – | – | – | + | + | + | + | – | – | + |
| *bla* CTX-M | – | – | – | – | – | – | – | – | – | – | – | – |
| September (spring) |  |  |  |  |  |  |  |  |  |  |  |  |
| *uidA* | + | + | + | + | + | + | + | + | + | + | + | + |
| *stx1* | – | + | – | + | – | – | – | – | + | – | + | + |
| *stx2* | – | + | – | + | – | + | + | – | – | + | – | + |
| *eae* | – | + | + | + | – | + | + | + | + | + | + | + |
| *wzy* 026 | – | – | – | – | – | – | – | – | + | – | – | – |
| *rfbE* O157 | – | – | – | – | – | + | – | – | – | + | + | + |
| *bla* CTX-M | – | – | – | + | – | – | – | – | + | – | + | + |

+ Gene detected
– Gene not detected

In autumn, the genes associated with pathogenic STEC O157 (i.e., *stx1, stx2, eae* and *rfbE*) were present at 33% of sample sites. All three virulence and both serogroup genes were detected in Rangitata River water and Ashley River sediment samples from below intensive dairy. Additionally, the *stx1, eae*, and *rfbE* genes were present in water from the site below intensive dairy on the Ashley River. No other virulence or serogroup genes were detected in the autumn samples.

In spring *stx1, stx2* and *eae* genes were detected in all rivers. These three STEC virulence genes were detected in sediment samples from five of the six sites (i.e., all except the Ashley below intensive dairy) but only from below intensive dairy in the water samples; only the *eae* gene was detected in water samples from above intensive dairy. In spring samples, the gene associated with the O157 serogroup (*rfbE*) was detected four times more often than that associated with the O26 serogroup (*wzy*) and the *bla* CTX-M gene was more frequent below intensive dairy. In three samples *stx1, stx2* and *eae* were detected together; in one of these the gene associated with the O157 serogroup was present but the genes associated with O157 and O26 were not detected in the remaining two. The *stx1* and *eae* genes were detected in two water samples from below intensive dairy whereas *stx2* and *eae* were detected in three sediment samples, two from below and one above intensive dairy.

## Discussion

The presence of human pathogenic STEC in New Zealand waterways is poorly documented. In part, this is because STEC strains have diverse metabolic requirements; since strains may co-occur, identifying individual organisms present in a sample would require multiple culture methods (Kerangart et al. 2018; Possé et al. 2007). In studies on STEC strains from ruminant faeces and environmental samples the microbial communities are generally enriched prior to DNA-based testing (Browne et al. 2018; Irshad et al. 2016; Jaros et al. 2013). The potential impacts of such enrichment on the microbial communities and on inferences concerning the detection of zoonotic disease are difficult to quantify. In the present study, we tested samples for genes associated with virulence and antibiotic resistance in STEC without an initial enrichment step.

In our analyses the genetic markers indicative of *E. coli* (*uidA*) and STEC virulence (*stx1, stx2* and *eae*) genes were detected in all three of the rivers sampled. That all samples tested positive for the *uidA* gene is consistent with *E. coli* being isolated at all six sites, from both substrates, and at both sampling times using traditional plating. Using relatively small sample volumes we detected genes associated with both human pathogenic STEC and antibiotic resistance without first enriching the microbial communities. Although all the STEC virulence and O serotype genes were detected in all rivers, and in some cases all these components were detected from a single sample, their presence varied by location, substrate, and season. These results suggest that explanations for the distribution of these genes across the landscape are likely to be complex and involve a range of factors.

*Escherichia coli* levels were higher and STEC associated genes more commonly detected at sites below intensive dairy. For example, the *stx1* gene was detected in six of 12 (50%) samples, and at least once at every site, below intensive dairy but in only two of 12 (17%) samples taken above intensive dairy. These results are consistent with previous reports suggesting that agricultural effluent is a primary source of faecal contamination in New Zealand waterways (Gluckman 2017). Studies have shown that faecal bacteria are transferred from pastures to waterways via run-off; carried either directly by the flow or indirectly as a result of adsorption to soil particles (Byappanahalli and Ishii 2011; Muirhead et al. 2004; Palmateer et al. 1993). However, studies have suggested that paddock-feeding waterfowl may also transmit ruminant hosted faecal bacteria when environmental contamination is high (Yang et al. 2019; Zou et al. 2019). In the current study we sampled adjacent to intensive dairy operations but expected microbial communities to vary along each river due to the diversity of land uses within each catchment. Microbial communities are likely to be most strongly influenced by adjoining land use but also impacted by inputs from upstream of the sampling location. Moreover, microbial communities reflect the prevailing environmental conditions and primary land use within the catchment. For example, the *stx1* and *stx2* genes were only detected at above intensive dairy sites during spring. One explanation for this observation may be inputs from smaller farms upstream of the reach bordered by the high densities of intensive dairying.

The STEC associated genes occurred at greater frequency in the spring sampling than the autumn sampling. Specifically, they were detected in eight of 12 samples (66%) in the spring and in three of 12 samples (25%) in the autumn. This result reflects a combination of increased detection below intensive dairy (i.e., five in September and three in May) as well as detection above intensive dairy in the spring. Increased detection of STEC and antibiotic resistance during the spring is likely a response to several factors. Spring calving may increase the load of pathogenic bacteria on land neighbouring these rivers. As calves have poorly developed intestinal biomes, are stressed by weaning, or are removed from their mother prior to receiving colostrum they are prone to colonization by and heavy shedding of bacteria, including STEC (Browne et al. 2018). In addition to the increased faecal loading on pastures, spring rainfall patterns may lead to higher faecal inputs reaching rivers. Other possible explanations for fewer detections in the autumn include lower base flow rates or drying out of rivers during the summer (e.g., the Selwyn), removal of stock from flood prone areas for winter grazing, and geomorphology that encourages adsorption of bacteria to sediments (Reddy et al. 1981). That STEC associated genes were more frequently detected during the spring is in contrast to the seasonal cycle of human STEC disease cases in New Zealand. Although there is a peak in notified human cases during spring, the autumn peak is typically larger. For example, in 2017 there were almost twice as many cases of STEC in autumn than during spring (Health and Environment Group ESR 2019). Moreover, although rates of notified human cases of STEC do differ between regions the differences are not always consistent with the prevalence of dairy farming. For example, in 2017 the Auckland and Waikato districts reported 7.4 and 8.6 cases per 100,000 individuals despite dairy farming being far more common in the Waikato region (Health and Environment Group ESR 2019). Taken together these observations suggest that livestock are unlikely to be the sole source of human STEC infections in New Zealand and that we need to be cautious when interpreting data on the distribution of STEC, or the genetic components linked to it, in the context of public health.

In all three of the sampled rivers *E. coli* levels were consistently higher in sediments than in the water column. The STEC associated genes were also more commonly detected in sediment samples. These results are consistent with previous studies that indicate aquatic sediments may act as a store for *E. coli* (Muirhead et al. 2004; Perkins et al. 2014; Wilkinson et al. 1995). Given that such stores may persist for months or years (Anderson et al. 2005; Garzio-Hadzick et al. 2010; Gerba and McLeod 1976) and that suspended sediments increase *E. coli* levels in the water column (Davies-Colley et al. 2018; Weiskerger and Whitman 2018) this finding has potentially important implications for water monitoring. In New Zealand, water quality testing by local government agencies is currently restricted to the water column. Although recreational use of waterways is discouraged when levels of suspended sediment are high (e.g., following precipitation) (Davies-Colley et al. 2018), disturbance of sediments by recreational users is not generally considered. Further work is needed to quantify the pathogens associated with the localised mixing of sediment into the water column by recreational users and to determine whether zoonoses are being underestimated by the sampling of a single substrate.

## Conclusion

Microbial culturing is not an efficient tool for monitoring STEC in recreational waterways. In part this can be attributed to the metabolic diversity of STEC and the apparent lack of a relationship between faecal indicator bacteria (e.g., *E. coli* counts) and the presence of genes associated with STEC. Understanding this relationship would require more intensive sampling over a broader geographic range. In most cases monitoring is conducted on samples retrieved from the water column. Such samples may not accurately reflect the microbial community that recreational users of the waterway may be exposed to. Our results suggest sediments may act as an important STEC reservoir and resuspension of these sediments by waterway users could potentially increase exposure to STEC.

There is a growing appreciation of new technologies that enable the presence and persistence of zoonoses to be monitored without the need for microbial culturing (Byappanahalli and Ishii 2011; Rose et al. 2001). This represents a fundamental change in our approach to microbial monitoring allowing us to take a holistic view of the riverine environment and improving our ability to ensure environmental, animal, and human health. While small, this study is the first step towards understanding zoonoses in New Zealand waterways at a time when global health is under the microscope.

## Supporting information

Author permissions

CFI form

## Acknowledgements

We thank Martin Taylor (Fish and Game New Zealand) for funding this study and providing staff to collect samples.

