## Supplementary material for "Detecting Genes Associated with Pathogenicity and Antimicrobial Resistance in Three New Zealand Waterways": Author permissions

Meredith T. Davis, Anne C. Midwinter, Russell G. Death, Richard Cosgrove, and Richard C. Winkworth the authors of “Detecting Genes Associated with Pathogenicity and Antimicrobial Resistance in Three New Zealand Waterways.” confirm that all authors have read the manuscript, agree that it is ready for submission and accept responsibility for its contents.
