## Supplementary material for "Detecting Genes Associated with Pathogenicity and Antimicrobial Resistance in Three New Zealand Waterways": CFI form

ehp Environmental Health Perspectives

### Competing Financial Interests Declaration

*Environmental Health Perspectives* maintains that authors are accountable for the articles submitted to the journal and requires authors to declare actual or potential competing financial interests that might be perceived as influencing the results or interpretation of a reported study. As indicated in the Instructions to Authors:

Authors must disclose actual and/or potential competing financial interests, including but not limited to grant support; employment (past, present, or firm offer of future); patents (pending or applied); payment for expert witness or testimony; personal financial interests by the authors, immediate family members, or institutional affiliations that may gain or lose financially through publication of the article; and forms of compensation, including travel funding, consultancies, board positions, patent and royalty arrangements, stock shares, or bonds. Diversified mutual funds or investment trusts do not constitute a competing financial interest. Authors employed by a for-profit, nonprofit, foundation, or advocacy group must also disclose employment. Authors should carefully examine the wording of documents such as grants and contracts to determine whether there might be an actual or potential competing interest.

All actual or potential competing interests occurring during the last 3 years should be reported. As a general rule, all consultants and contractors must indicate a potential competing financial interest and provide the name of their employer and a description of any specific potential conflicts that exist as a consequence of their employment.

Corresponding authors are required to submit this form declaring actual or potential competing financial interests on behalf of all authors involved. Failure to declare a competing financial interest could result in a ban on publication for 3 years and a retraction of the article. Disclosure of actual or potential competing interests does not imply that the information in the article is questionable or that conclusions are biased. However, the corresponding author must certify that the authors' freedom to design, conduct, interpret, and publish research is not compromised by any controlling sponsor as a condition of review and publication (see below).

Detecting Genes Associated with Pathogenicity and Antimicrobial Resistance in Three New Zealand Waterways.

Manuscript number (if available) and title

Meredith T. Davis, Anne C. Midwinter, Russell G. Death, Richard Cosgrove, and Richard C. Winkworth

Authors

Please check one of the following:

- [ ] One or more authors declare they have an actual or potential competing financial interest (see above for examples). Provide details below.
- [x] All authors declare they have no actual or potential competing financial interest.

Specify the actual or potential competing financial interests, including the name of entity, who is involved, and current interest.

I am unsure if this is a “ competing financial interest ” as none of the research would result in financial gain or loss, but Richard Cosgrove works for Fish and Game New Zealand who paid for the sample processing.

Are there any other relevant disclosures that should be brought to the editors' attention?

On behalf of the authors of this paper, I hereby certify that all actual or potential competing financial interests have been declared and that the authors' freedom to design, conduct, interpret, and publish research is not compromised by any controlling sponsor as a condition of review and publication.

| | |
|---|---|
| [signature] | April 17, 2020 |
| Signature | Date |
| Meredith T. Davis | Massey University |
| Print Name | Affiliation |
